# Growth Cost and Transport Efficiency Tradeoffs Define Root System Optimization Across Varying Developmental Stages and Environments in Arabidopsis

**DOI:** 10.1101/2025.07.25.666579

**Authors:** Kian Faizi, Preyanka Mehta, Amy Maida, Taylor Humphreys, Elizabeth Berrigan, Leo McKee-Reid, Robbie McCorkell, Arush Tagade, Jessica Rumbelow, Julia Showalter, Lukas Brent, Clementine Coroenne, Audrey Rigaud, Arjun Chandrasekhar, Saket Navlakha, Antoine Martin, Christophe Pradal, Sanghwa Lee, Wolfgang Busch, Matthieu Pierre Platre

## Abstract

Root system architecture (RSA) is central to plant adaptation and fitness, yet the design principles and regulatory mechanisms connecting RSA to environmental adaptation are not well understood. We developed Ariadne, a semi-automated software for quantifying cost-efficiency tradeoffs of RSA by mapping root networks onto a Pareto-optimality framework, which describes the balance between resource transport efficiency and construction cost. Applying Ariadne to *Arabidopsis thaliana*, we found that root architectures consistently assume Pareto-optimal forms across developmental stages, genotypes, and environmental conditions. Using the Discovery Engine, an engine that combines machine learning together with interpretability techniques, we found developmental stage, the *hy5/chl1-5* genotype, and manganese availability as important determinants of the cost-efficiency tradeoff, with manganese exerting a unique influence not observed for other nutrients. These results reveal that RSA plasticity is genetically constrained to cost-efficiency optimal configurations and that developmental and environmental factors shift RSA on the pareto front, with manganese acting as a strong modulator of the transport efficiency and construction cost balance.

## INTRODUCTION

The root system is a critical organ to ensure the anchoring, water and nutrients uptake, and photo-assimilate transport, necessary for plant survival. To maintain these functions when exposed to environmental cues, the root system alters its 3-dimensional organization, defined as the root system architecture (RSA)^1^. Depending on the environmental changes, a single genotype can display a variety of RSAs to adapt accordingly, highlighting the plasticity of the root system^1^. RSA plasticity is a key determinant of plant fitness and adaptation to adverse conditions. RSA is also highly relevant in the context of climate change, as elucidating the basis of RSA plasticity in an integrative manner is crucial for uncovering and enhancing plant resilience. Moreover, the root system represents a powerful lever to potentially sequester carbon underground to decrease atmospheric CO_2_ levels and mitigate climate change. This could be achieved by increasing one of the most impactful root traits for RSA, the root growth rate, which would maximize the size of the root system^2^. In addition, deeper RSA would benefit to mitigate climate change by sequestering carbon for a longer time-scale deeper in the soil^2^. It was estimated that substantially increasing the root depth and root mass of major crops could be the fastest pathway to achieving gigaton-scale CO₂ removal from the atmosphere^3^. Therefore, understanding how to develop crops that are resilient and maximize carbon storage would be a major step toward ensuring food security and mitigating climate change. This goal could be achieved by uncovering the constraints on RSA plasticity and the genetic mechanisms that control it.

Root architectural plasticity can be assessed by changes of morphological (e.g. length), geometrical (e.g. diameter), dynamical (e.g. growth rate), and topological descriptors^4^. During the last decades, tremendous efforts were made to study those first three aspects due to their relatively easy assessment. Nonetheless, the analysis of RSA topology, which consists of studying the arrangement of nodes and links of a network to assess its efficiency and functions, remains underexplored. Yet, these descriptors are particularly relevant because they not only integrate several RSA traits but also, they allow to directly assess the efficiency and functions of the RSA compared to the other non-topological descriptors.

The functions of the root system are, i) to transport nutrients, water and photo-assimilates and ii) to forage the soil for water and nutrients through growth and to promote anchoring of the plant. Therefore, the root system has to achieve two major objectives, transport, and growth. In network theory these objectives are considered two competitive objectives: efficiency (transport) and cost (growth)^5^. In the ideal case, the network needs to display an architecture that transports resources efficiently throughout the network (efficiency) while minimizing its growth (cost). Importantly, the improvement of one objective impacts the other one negatively, therefore always presenting a cost-efficiency tradeoff. The trade-off happens because maximizing transport efficiency usually involves investing in longer or denser root systems, which increases construction and maintenance costs. Conversely, minimizing cost means cutting back on network size or density, which can hamper efficient transport across the root system. For a wide range of biological or human made transport networks, such as railways, their architectures were shown to follow this principle revealing its universality^5–7^. Recently, the cost-efficiency tradeoff of *Solanum pimpinellifolium* RSA was tested for plants grown *in vitro* using a graph theoretic algorithm assessing Pareto optimality. In this algorithm the root system is considered as a transport network allowing to analyze the root system topological information. It processes the positioning of the nodes corresponding to the hypocotyl, the root tips, and junctions between two roots but also the links corresponding to the root material that connects the nodes. Based on these RSA topological information and by estimation of the *bona fide* cost and efficiency values of the RSA, they revealed that the RSA of *Solanum pimpinellifolium* follows a Pareto optimal cost-efficiency tradeoff.

In biological transport network analysis, a long-stated hypothesis is that through evolution, natural selection pushed architectures towards the Pareto optimality since most transport networks of living organisms display pareto optimal solutions^5–7^. However, it hasn’t been addressed whether and to which extent plants follow the Pareto optimal cost-efficiency tradeoff across different growth conditions, developmental stages and stresses. If broadly observed, this would imply that the root system is developmentally constrained by a genetic framework to constrain architectures to configurations that remain Pareto optimal in any condition while the specific cost-efficiency tradeoff may vary.

In this study, we developed a semi-automated tool, Ariadne, to universally analyze RSA across developmental stages and conditions. Our results show that root architectures consistently follow Pareto-optimal design principles, with cost-efficiency tradeoffs shaped by developmental and environmental factors, such as water and nutrient availability. Together, these findings suggest that RSA plasticity operates within a genetically constrained framework and provide an integrative approach to link root system design with resilience and carbon storage potential.

## RESULTS

### A semi-automated software to determine the Pareto optimal cost-efficiency of the root system network

To evaluate the Pareto optimality cost-efficiency tradeoff of the RSA in a wide range of plant species and conditions, we developed a user-friendly and versatile software named Ariadne. This software allows the processing of images of root systems in 2 dimensions of a variety of image formats (tif, jpeg, png, pdf, gif,…, Figure 1a). The root system is traced by the user through the graphical user interface of the software (Figure 1b). This allows one to generate a point cloud data set and store it in a .json file which represents the RSA tree like structure (Figure 1b). This file is then automatically processed by the Pareto algorithm to determine i) whether the RSA is pareto optimal and ii) its cost-efficiency tradeoff. We made this process compatible with the root system ML (RSML) format to make it available for a wider range of users^8^.

**Figure 1:**
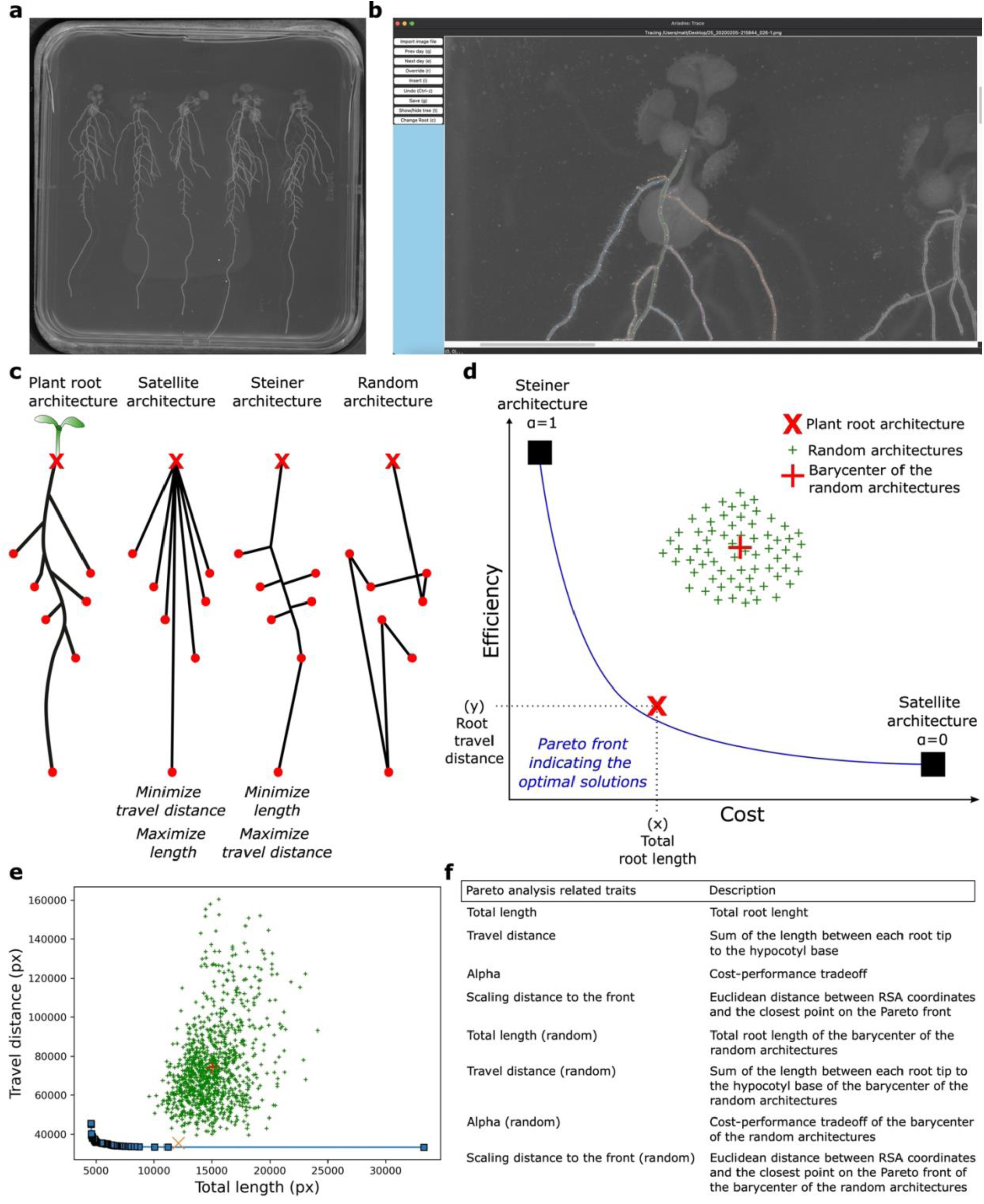
Presentation of the cost-efficiency Pareto optimality concept and the software Ariadne. **a**, Representative image of plants grown *in vitro* in mock conditions in plates for 14 days. **b**, Representative image of the interface of Ariadne. **c**, Schematic representation of, the plant root system, the Satellite, the Steiner, and the random architectures, from left to right. The red crosses depict the hypocotyl bases and the red dots the root tips. **d**, Graphical representation of the so-called Pareto front to determine the optimality and the cost-performance tradeoff of the architecture of the root system. **e**, Output file of the Ariadne software representing the pareto front. **f**, Table depicting the root system architectural traits used to determine the Pareto optimality cost-performance tradeoff.

The algorithm draws the two most extreme virtual architectures from the positions of the nodes corresponding to the base of the hypocotyl and the root tips (Figure 1c). We defined the base of the hypocotyl as the input point because it represents the convergence of the shoot and root systems, serving as a central hub through which all transported molecules pass. As additional input points, we selected the root tips, since root growth originates there through meristematic activity and elongation, supported by the transport of photosynthetic products via the phloem. In addition, approximately 1.5 mm above the root tip, within the maturation zone, root hairs begin to emerge and actively absorb water and nutrients. These root hairs are short-lived, typically lasting only from days to a couple of weeks, and play a key role in nutrient uptake compared to other regions of the mature root^9,10^. One model RSA represents the minimum travel distance between each root tip and the base of the hypocotyl creating straight links between each root tip and the hypocotyl. This is called the satellite tree (Figure 1c)^6^. The other model RSA represents the minimal building cost of the architecture connecting all the nodes with minimal total length while allowing the addition of extra points that serve as branching points. This is called the Steiner tree (Figure 1c)^6^. According to graph transport network theory, which describes cost and transport efficiency as competing objectives, the Satellite tree exhibits high transport efficiency at a higher cost, while the Steiner tree shows lower cost but reduced transport efficiency. (Figure 1c). These two extreme architectures serve as an anchor point to generate all Pareto optimal solutions displaying intermediate optimal cost-efficiency tradeoffs (Figure 1d). They are illustrated on a graph by the so-called Pareto front and each position on this front is assigned an alpha value that ranges between 0 (Satellite tree) and 1 (Steiner tree, Figure 1f)^6^. To produce these solutions that lie on the Pareto front between Satellite and Steiner, we use the most straightforward joint objective: a linear combination of the two separate objectives.

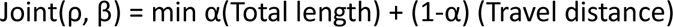

where 0≤ *α* ≤1. When *α* =0, the optimal solution of the combined objective corresponds to the Satellite. When *α* =1, the optimal solution matches the Steiner tree. The total length is represented by the entire length of the network while the travel distance is represented by the sum of the length from the hypocotyl to each root tip. ρ being a set of points in the 2D Euclidean space, with the point ρ0 represents the base of the hypocotyl, while the other n points indicate the root tips. β corresponds to undirected branches or edges that exist between ρ0 and the root tips. The joint optimization problem is NP-hard^11^.

Having this framework, the measured plant RSA is projected on the Pareto front graph using its bona fide cost (x, total root length) and transport efficiency (y, travel distance) values (Figure 1d). Then, we measured the distance between the position of the RSA to the Pareto front by determining the Euclidean distance between the RSA to the front (hereafter referred as the scaling distance to the front). We named this distance the Pareto optimality value. This procedure allows to precisely position the RSA relative to the Pareto front and thus enables the quantification of the alpha value of any RSA indicating the cost-efficiency tradeoff (Figure 1d) as well as the levels of “optimality”. To assess the likelihood of any given plant RSA to be Pareto optimal, thousands of randomly generated spanning trees (green crosses on the graph) are generated from the input nodes (Figure 1d). The barycenter of these random trees is calculated to obtain one representative random tree for which the distance from the Pareto front is calculated and compared to that of the plant RSA (Figure 1d). Altogether, these calculations allow to generate a graph for each RSA (Figure 1e), from which quantitative data are extracted to evaluate the RSA’s Pareto optimality and its associated cost-efficiency trade-off (Figure 1f). Additionally, the software incorporates various geometrical, morphological, and, when using time-series images, dynamic features, enabling integrative analysis of root system architecture (RSA) (Figure 1f and Table S1). The tool is publicly available on the Pypi platform (https://pypi.org/project/ariadne-roots/). To conclude, Ariadne offers a semi-automated software to evaluate the Pareto optimality and cost-efficiency tradeoff along with several canonical RSA traits for all types of 2D images.

### The *Arabidopsis thaliana* root system represents a cost-efficiency Pareto optimal solution

Using the Ariadne software, we set out to analyze the Pareto optimality of the RSA of the model plant *Arabidopsis thaliana* (Arabidopsis). We first analyzed the RSA of Arabidopsis from 2D drawings of excavated root systems from the “Wurzelatlas” that was made publicly available by Wageningen University^12,13^. We found that the Arabidopsis root system grown under natural conditions in soil lies closer to the Pareto front than any of our randomly generated architectures with scaling distances to the front of 1.16 and 7.99, respectively (Figure 2a). This shows that this plant species follows the universal design principle of the Pareto optimal cost-efficiency tradeoff. Moreover, we observed that the alpha value representing the cost-efficiency tradeoff was 0.03 (Figure 2a), showing that Arabidopsis in natural conditions tends to display an architecture that favors the transport of nutrients and water at the expense of the growth cost.

**Figure 2:**
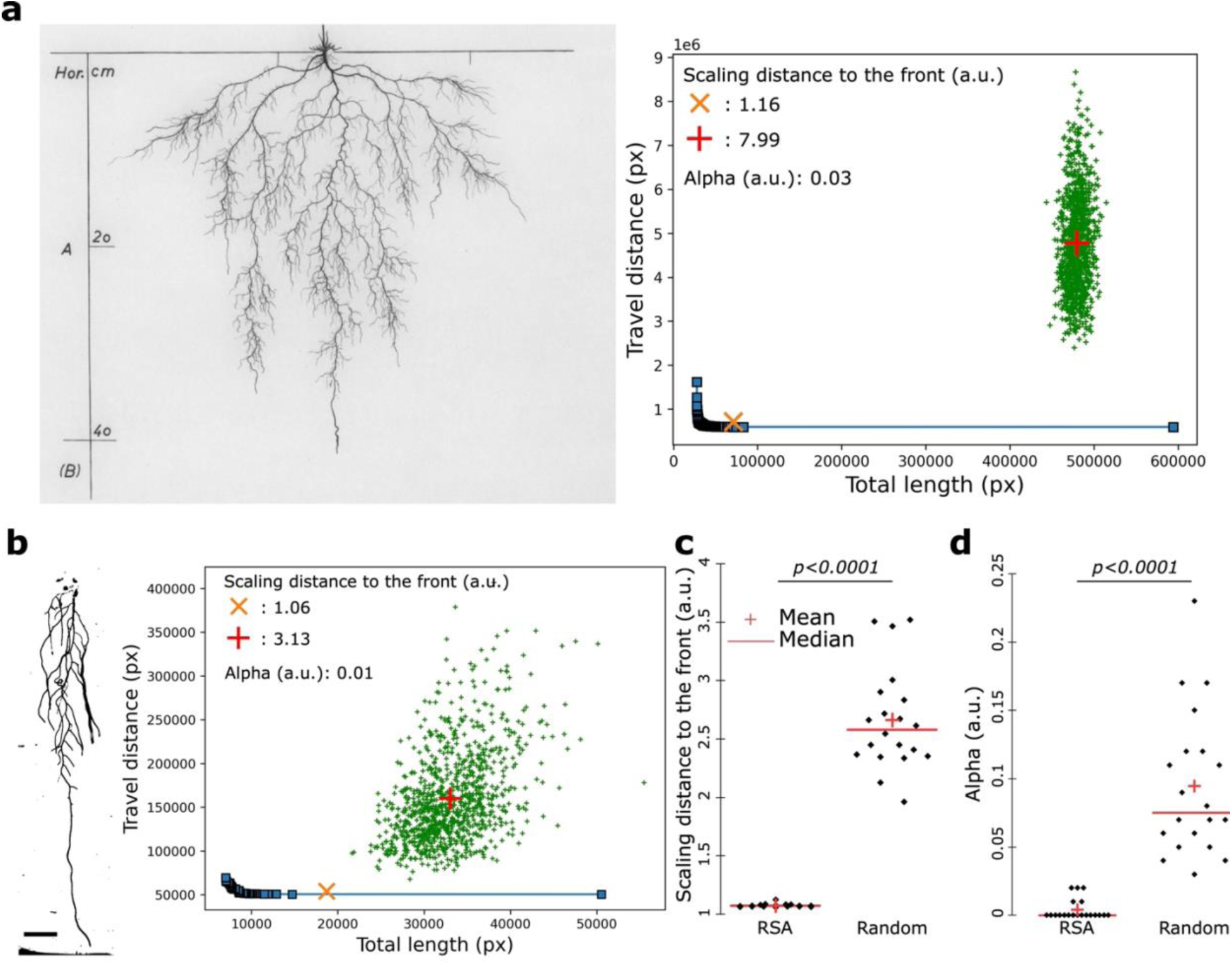
The root systems of Arabidopsis thaliana is Pareto optimal for the cost-efficiency in natural and *in vitro* conditions. **a**, 2D scan of the *Arabidopsis thaliana* root system in natural conditions (left) and the corresponding graphical representation of the cost-performance Pareto optimality (right). **b-d** 2D scan of the *Arabidopsis thaliana* root systemin grown *in vitro* (left, b) and the corresponding graphical representation of the cost-performance Pareto optimality (right, b) with the corresponding quantification of the scaling distance to the front (**c**) and alpha value (**d**), [Two-ways Mann-Whitney test, p=0.05]. the red crosses depict the mean and the red bars the median. Scale bar, 1cm.

We then assessed the changes of the RSA cost-efficiency tradeoff in other conditions. To do so, we evaluated the Pareto optimality of Arabidopsis grown *in vitro* on vertical plates for several weeks. This experiment showed that under these conditions, Arabidopsis RSA recapitulated the patterns we previously observed in natural environments (Figure 2b). It displayed a distance to the front of 1.06 which is closer to the front compared to the random one (3.13) and an alpha value indicating the cost-efficiency tradeoff of 0.01 showing that like in soil, Arabidopsis root systems on agar plates assume an architecture that favors transport over growth cost (Figure 2b). Moreover, extending this calculation to a large number of plants revealed a highly significant difference between the RSA from real plants and random architectures (Figure 2c). While random RSAs showed a scaling difference of approximately 2.6, Arabidopsis root systems consistently exhibited a scaling difference close to 1 (Figure 2c). Taken together, this set of experiments demonstrated that Arabidopsis root systems follow a Pareto optimal solution for the cost-efficiency tradeoff independently of their growth conditions. In addition, this extended analysis confirmed also that Arabidopsis root systems assume architectures that can transport molecules efficiently rather than growing at low cost indicated by the alpha value of 0.004, which is several magnitudes less than randomly generated RSAs, which show a mean alpha value of almost 0.1 (Figure 2d). In conclusion, Arabidopsis root systems consistently adopt a Pareto-optimal configuration that strongly favors transport efficiency over growth cost, even across markedly different growth conditions (for example, in sandy clay soil in Klagenfurt (Austria) in 1960 and on agar plates in La Jolla, California (USA) in 2022).

### The Pareto optimal cost-efficiency tradeoff of the root system is developmentally controlled

We demonstrated that Arabidopsis RSA grown under natural and agar conditions assume Pareto optimality for the cost-efficiency tradeoff (Figure 2). However, we noticed that the Pareto optimal and the alpha values were different between the Arabidopsis grown under natural and agar conditions with a pareto optimality value of 1.16 and 1.06 and alpha value of 0.03 and 0.01, respectively (Figure 2). We decided to explore the nature of these variations. First, we hypothesized that the developmental stage could account for these differences, as the Arabidopsis plant grown under natural conditions was an adult plant, presumably several months old and at the flowering stage, with a root depth of 42 cm. In contrast, the plants on agar plates were grown for 17 days and imaged before reaching the flowering stage, displaying root depths of less than 12 cm (Figure 2). To assess the effect of developmental stage on the cost-efficiency tradeoff, we performed time-lapse imaging, capturing an image each day beginning with the appearance of the first lateral root, which occurred approximately 7 to 9 days after germination under our conditions until plants reached the bottom of the plate which is about 17 to 18 days after sowing. In line with our expectations, we observed changes in both measures over the different developmental stages, albeit with stronger effects on the alpha value rather than the optimality (Figure 3a,c). In addition, even though we observed that the RSA is finding different optimal solutions during development, all of them were representing a higher level of optimality than random architectures according to the scaling distance to the front between the RSA and random architectures (Figure 3d). Altogether, this suggests that over development the Pareto optimality value and cost-efficiency tradeoff of the RSA change and might be genetically encoded.

**Figure 3:**
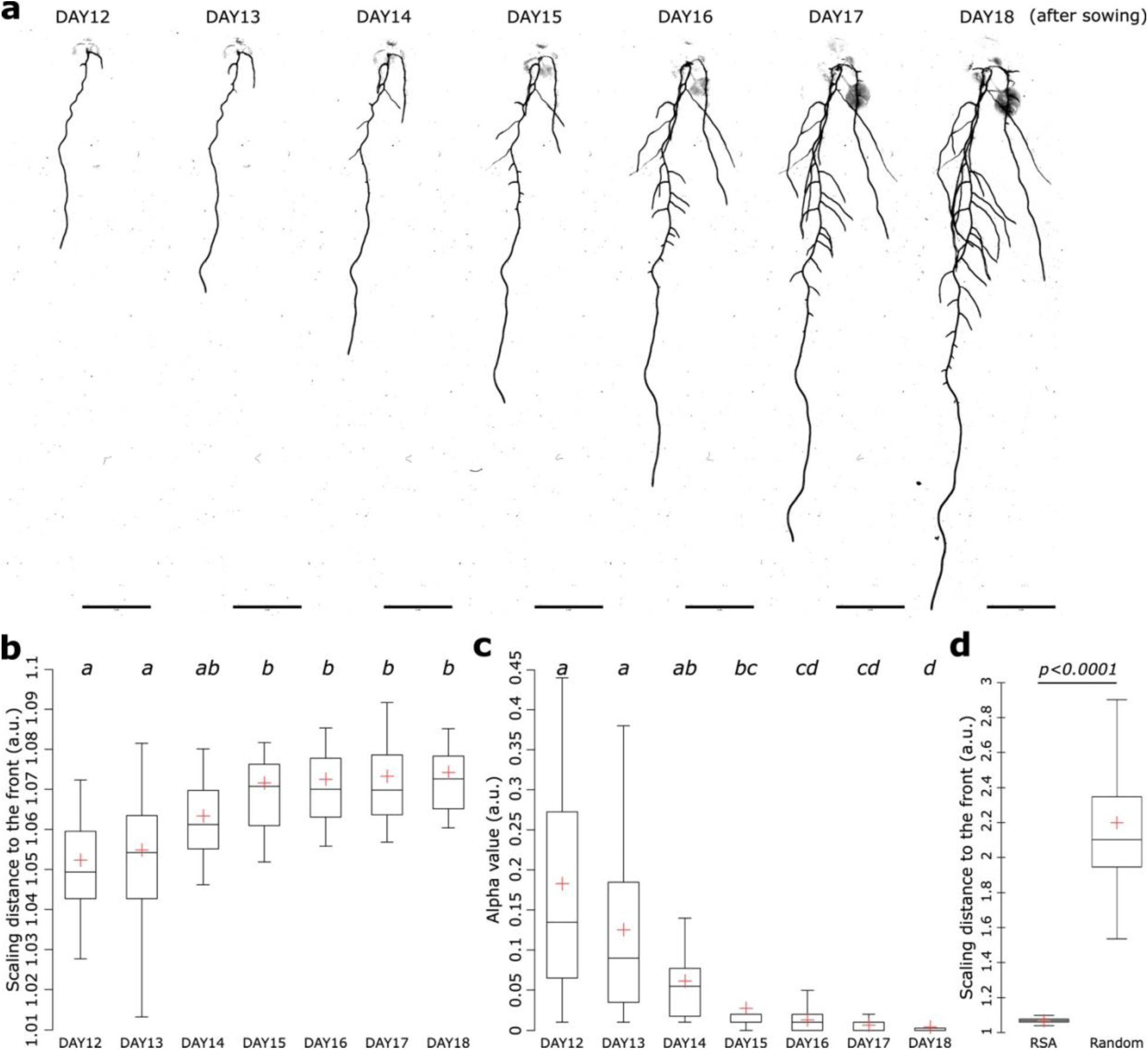
The Pareto optimal cost-efficiency tradeoff varies during development in *Arabidopsis thaliana*. **a-d**, Images of the developmental of the root system from day 7 to 13 (**a**) and the related quantification of the scaling distance to the front (**b**) [two-way ANOVA, Tukey HSD test, letters indicate statistical differences (p < 0.05)], alpha value (**c**) [two-way Kruskal-Wallis coupled with post hoc Steel-Dwass-Critchlow-Fligner procedure was performed, letters indicate statistical differences (p < 0.05)] and the scaling distance to the front for all days of RSA and random architectures (**d**) [Two-ways Student t-test (p=0.05)]. The red crosses depict the mean. Scale bar, 1cm.

### Osmotic stress-induced water deficit alters the root Pareto optimality and the cost-efficiency tradeoff

We found that developmental stages might explain the differences observed for the Pareto optimality and cost-efficiency tradeoff values between plants grown under natural conditions and agar conditions (Figure 3). However, these differences might also be explained by environmental factors to which RSA is known to respond strongly. Water availability has one of the most profound effects on RSA. We therefore set out to test the impact of water availability on the Pareto optimality and cost-efficiency tradeoffs. For this, we first subjected plants to osmotic stress by applying sorbitol at 150mM to decrease plant water availability, which is known to drastically affect the RSA^14,15^. As previously reported, this condition significantly decreased the primary root length, confirming our experimental set up (Figure S1a). When analyzing the scaling distance to the Pareto front and alpha value, we observed a decrease and an increase relative to the mock condition, respectively (Figure 4). The most notable change was in the cost-efficiency tradeoff value which increased from 0.02 under control conditions to 0.1 under sorbitol treatment (a fivefold difference). This indicates that roots invest relatively less building material under drought-like conditions compared to non-drought conditions, while achieving an even slightly better tradeoff optimality under water limitation (even though the distance to the Pareto front only varied slightly, decreasing from 1.07 to 1.04). Although the optimal solutions were different between mock and Sorbitol conditions in both cases, they were displaying higher Pareto optimality than random architectures according to the scaling distance to the front (Figure S1b). This indicates that both, optimality on the cost-efficiency spectrum and distance to the optimum change upon exposure to stresses.

**Figure 4:**
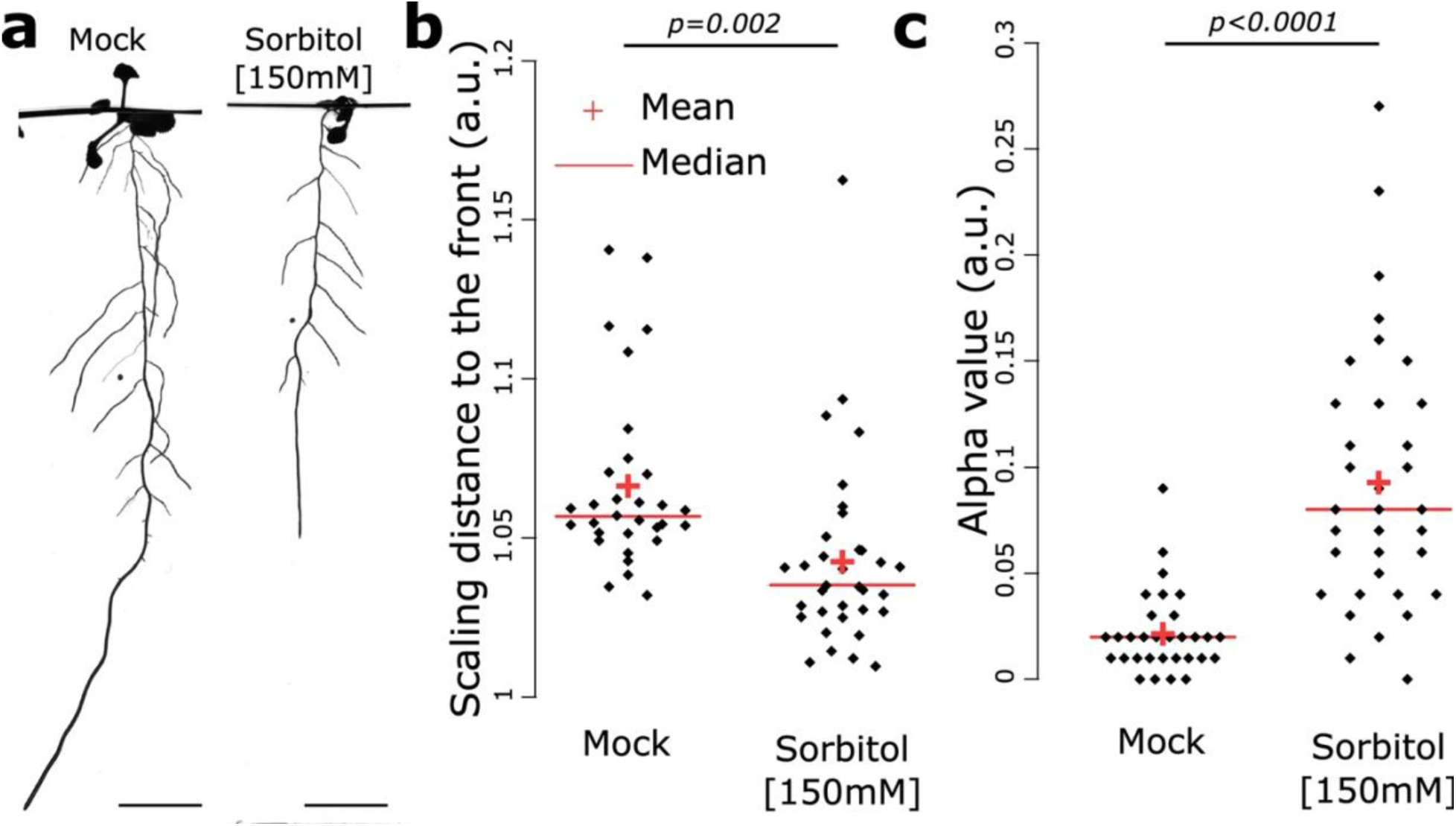
The Pareto optimality and cost-efficiency tradeoff is altered upon water deficit. **a-c**, 2D scan of the *Arabidopsis thaliana* root system under mock and sorbitol 150mM (**a**) and the corresponding quantification of the scaling distance to the front (**b**), and alpha value (**c**). [Two-ways Student t-test (p=0.05). The red crosses depict the mean and the red bars the median. Scale bar, 1cm.

### Comprehensive assessment of RSA plasticity across developmental stages, nutrient regimes and genotypes using interpretable machine learning via the Discovery Engine

After demonstrating the impact of water-limited conditions, we next sought to identify additional environmental factors that might influence the cost-efficiency tradeoff. To do so, we leveraged the RSA plasticity in various conditions relating to nutrient availability and status. For this, we used 27 environmental conditions and mutations affecting RSA^16^: *elongated hypocotyl 5* (*hy5*), *chlorina 1-5* (*chl1-5*), *hy5;chl1-5* double mutant and *brassinosteroids insensitive 1-like 3* (*brl3-2*) and wild type (WT) as a control. *HY5* regulates root responses to environmental cues^17,18^, *CHL1* is a dual-affinity nitrate transporter also involved in auxin transport^19,20^, and *BRL3* is a brassinosteroids receptor implicated in hydrotropism and stress responses, though its role in RSA is less clear^21–24^. Mutants were grown on agar plates under control and nutrient-deficient conditions (low nitrate, phosphate, or both), and WT and *brl3-2* were exposed to gradients of N, Mn, Mg, and P concentrations (Table S2). RSA traits were monitored daily from the emergence of the first lateral root until the primary root reached the plate bottom, covering days 7–21 after sowing. This comprehensive setup spanning developmental stages, genotypes, and nutrient conditions yielded about 180 000 RSA trait measurements on more than 10 000 plants and 18 RSA traits (Table S3).

To efficiently identify factors impacting the cost-efficiency tradeoff (alpha value), we used the Discovery Engine, a machine learning platform tailored for scientific discovery through the use of interpretable technics^25,26^. We reasoned that this approach would be particularly well-suited for our study, where the goal was to uncover RSA patterns across diverse genotypes, developmental stages, and environmental conditions. Unlike classical analytical methods such as PCA, correlation, or standard graph analysis that are limited to uncovering linear relationships and may overlook complex, nonlinear interactions, the Discovery Engine is capable of capturing intricate dependencies within high-dimensional, heterogeneous datasets like ours. It also enables rapid and automated exploration of large datasets that would otherwise be extremely laborious and time-consuming to analyze manually. Most importantly, this methodology is inherently unbiased, as it allows for the objective identification of patterns and relationships without preconceived assumptions or manual selection of features. Furthermore, interpretability tools not only highlight which factors are most influential but also reveal how these factors interact to drive observed outcomes. As a result, we reasoned that this methodology could enable the discovery of key regulators and mechanisms underlying RSA plasticity that would likely remain hidden using traditional analyses, providing deeper and more actionable biological insights. The discovery machine is divided into 4 critical steps (Figure S2).

1. <u>Data Ingestion</u>: This initial step involves automatic preprocessing of the trait data. Key operations include imputing missing values, removing duplicates, eliminating highly correlated columns, and managing both categorical and continuous variables to prepare the dataset for modeling^26^. In our case, we condensed the dataset from 180 000 to about 12 000 RSA measurements using the average of about 15 plants per condition, genotype, and day. In addition, we processed a one-hot encoding to transform the categorical data into a format where each category is represented as a binary vector.
2. <u>AutoML</u>: The preprocessed data is then modeled using a tailored AutoML component designed specifically for scientific discovery. Unlike general AutoML tools that emphasize transfer learning and lack interpretability, this system incorporates a variety of models which includes linear (e.g., linear regression), tree-based (e.g., XGBoost), kernel-based (e.g., SVM), and deep learning models (e.g., autoencoders)^27–29^. By supporting both simple and complex models, it aims to avoid overfitting and aids in the discovery of relevant data relations. For our dataset, we trained the model using a 70:30 train–test split, ensuring a balance between providing sufficient data for learning and retaining enough samples for reliable evaluation^30^.
3. <u>AutoInterp</u>: Once models are trained, a suite of interpretability techniques is applied to uncover data patterns. These include feature importance measures, identification of top representative training examples, generation of prototypical synthetic examples that maximally activate specific classes or regression values, and global counterfactuals that reveal minimal alterations switching class membership or output value which are crucial for understanding and exaggerating learned patterns.
4. A<u>utoEvaluate</u>: The interpretability outputs are analyzed to prioritize robust novel patterns while filtering out noise. This step involves grouping and ranking artefacts collectively, with the assistance of large language models (LLMs) to explain and contextualize results by referencing external knowledge sources like scientific literature. Finally, all extracted patterns undergo validation on the original dataset before being presented as reliable discoveries. A report is then generated based on the validated patterns.

### The Discovery Engine uncovers the complex interactions among multiple growth parameters that contribute to root plasticity

To validate the findings of the Discovery Engine, we first chose to focus on total root length, as it is a well-established trait that has been extensively studied and widely characterized in relation to genotype, nutrient availability, and developmental stage. The architecture of the Discovery Engine model for the target, total root length, was a fully connected feedforward neural network consisting of three linear layers. The first linear layer mapped 35 input features to 45 output features (i.e. hidden neurons), followed by a ReLU activation. The second layer reduced the dimensionality from 45 to 23 features, again followed by a ReLU activation. The final linear layer mapped the 23 features to a single output. After training, we found that the model displayed high performance according to the Mean Absolute Error (MAE), Root Mean Squared Error (RMSE) and R-squared (R²) scores for predicting this trait (Supplemental S1). We therefore assumed that the model could predict complex and non-linear relationships already identified in the literature for the total root length. The Discovery Engine identified that total root length peaked at day 20 and was lowest between days 8 and 11, consistent with time being the most relevant variable for root growth since root growth increases root length during plant development (Table S4, Figure S3a,b). However, we observed that although the dataset included RSA traits spanning days 7 to 21, the model did not accurately predict the minimum and maximum total root lengths corresponding to the start and end of this period. This discrepancy might be due to our analysis with Ariadne specifically focusing on the interval between the emergence of the first lateral root and the moment the primary root reached the bottom of the plate—events whose timing varied considerably across samples. As a result, the earliest and latest time points were underrepresented in the original dataset, which likely reduced the model’s ability to accurately predict these extreme values (Figure S3c). Discovery Engine then identified the conditions in which total root length was maximized within the entire dataset. This condition was plants grown 5 days in 11.4mM N then transferred to 0.275mM N at age 14–17 days, reflecting known root foraging under low N (Table S4)^16,31^. This finding was also compared with the full original dataset which confirmed the finding that total root length under 0.275mM of N is higher after several days (day 11 corresponding to 6 days after transfer to low N) in this condition compared to 11.4mM of N (Figure S3d). Similar to the previous observation the lack of data points in the original data set for the older days might explain the inability of the model to predict higher total root length for later days than 17 (Figure S3e). Nonetheless, the Discovery Engine was able to reveal that the root foraging response is temporally regulated. Conversely, the model identified that the condition with most decreased root length was occurring in *hy5;chl1-5* mutants at days 10–13. Again, we confirmed this by comparing to the entire dataset (Table S4 and Figure S3f). This aligns with prior findings of reduced primary root length for *hy5;chl1-5* mutant^17^. When further analyzing the original dataset, we observed that indeed, this genotype presented a lower total root length during this period (especially day 11 to 15) compared to all other genotypes grown in the same conditions (Figure S3g). This suggests that HY5-CHL1-5 module controls root growth processes over time possibly by its role in coordinating N-P nutritional responses^17^.

In summary, our approach combining large-scale phenotyping with interpretable machine learning enabled the unbiased identification of key genetic and environmental factors influencing root system architecture plasticity. The model not only recapitulated known root growth responses to nutrient availability and developmental timing but also was able to pinpoint specific periods during which root foraging and HY5-CHL1-5 axis was most critical in our conditions. These findings highlight the power of data-driven, interpretable methods to uncover nuanced, time-specific mechanisms underlying plant responses to the environment.

### Manganese levels and *HY5-CHL1-5* axis are major factors in controlling the root system cost-efficiency tradeoff

With Ariadne and the Discovery Engine, we established a pipeline that effectively identifies key factors influencing RSA traits. We then applied this to uncover factors influencing the cost-efficiency tradeoff, represented by the alpha value. The architecture of the model for the target, alpha, was a fully connected feedforward neural network composed of three linear layers. The first layer mapped 35 input features to 85 output features, followed by a ReLU activation. The second layer reduced the dimensionality from 85 to 43, again followed by a ReLU activation. The final linear layer mapped the 43 features to a single output. After training, the model showed relatively good performance in identifying factors underlying the alpha value, as indicated by medium high MAE, RMSE and R² (Supplemental S1). The interpretable machine learning model indicated that during days 8–12 alpha value is maximized and that during days 15 and 18 it is lower, which were confirmed in the original dataset with high significance (Table S4 and Figure S4a-b). This was also consistent with our previous results highlighting the importance of developmental stage on the alpha value (Figure 3). Like for total root length, due to the lack of data points for the earliest and latest time points, the model was not able to predict earlier or later days than 8 and 18 days, respectively (Figure S3c). In addition, the model pinpointed the genotype *hy5/chl1-5* to be a major factor explaining low alpha values in the dataset. We confirmed this in the data set (Figure S4c). This finding further supports the genetic control of the alpha value as previously suggested by our analysis of developmental stage (Figure 3). When then examined nutritional factors affecting the alpha value. The model pinpointed manganese (Mn) as most important determinant. Plants grown for 5 days at 0µM Mn and then transferred to 50µM Mn showed significant alpha minimization in the model and original dataset, (Table S4 and Figure S4d). We then set out to explore the impact of Mn concentrations on the alpha value. For this, we dissected the original dataset in detail. We analyzed 17-day-old wild-type plants that had been grown for 5 days in low-nutrient conditions (Mn 0 µM, N 0.11 µM, Mg 0 µM) and then had been transferred to high-nutrient conditions (Mn 50 µM, N 11.4 µM, Mg 750 µM) for 12 days. This developmental stage was selected because alpha values are lowest in older plants (Figure 3c, S3a-b). This Mn condition led to the lowest alpha values compared to all other conditions (Figure 5a). Given Mn’s similarity to Fe, we also compared plants that had been grown on 5 days in Fe-depleted medium (no iron with addition of the strong iron chelator FerroZine at 100µM) and then had been transferred to Fe-sufficient medium (75µM) for 12 days. Our results indicated that the profound effects on the alpha value are Mn specific (Figure 5a), suggesting that Mn availability directly modulates the cost-efficiency tradeoff. Testing eight Mn conditions, with plants first grown 5 days at 0µM or 50µM, then transferred to 0, 0.5, 10, or 50µM Mn, revealed a negative correlation between Mn levels and alpha (Figure 5b), a unique effect to Mn among all tested nutrient conditions (Figure S5). Thus, manganese levels directly fine-tune the trade-off between root building cost and transport efficiency. The insights generated by the Discovery Engine, together with their experimental validation, guided us in designing our next experiment. In Arabidopsis, the Mn homeostasis is regulated by the transporter *NATURAL RESISTANCE-ASSOCIATED MACROPHAGE PROTEIN 1 (NRAMP1)*^32^. In accordance, it was shown that *nramp1-1* mutant contains less Mn levels in the root *in vitro* in manganese sufficiency conditions^32^. Therefore, we set out to test the cost-efficiency tradeoff evaluating the alpha value in *nramp1-1* mutant compared to WT in Mn sufficient levels. To our surprise, we did not observe any difference compared to the WT (Figure 5d,e). In addition, *IRON REGULATED TRANSPORTER 1 (IRT1)* is another transporter able to transport Mn and decided to test the alpha value in plants defective for this gene using *irt1-1* mutant plant^33^. In this mutant, we could not detect any difference with the WT (Figure 5d,e). Altogether, these experiments show that the two Mn transporters, *NRAMP1* and *IRT1* do not impact the cost-efficiency tradeoff, yet, external manganese levels are crucial to fine tune this tradeoff.

**Figure 5:**
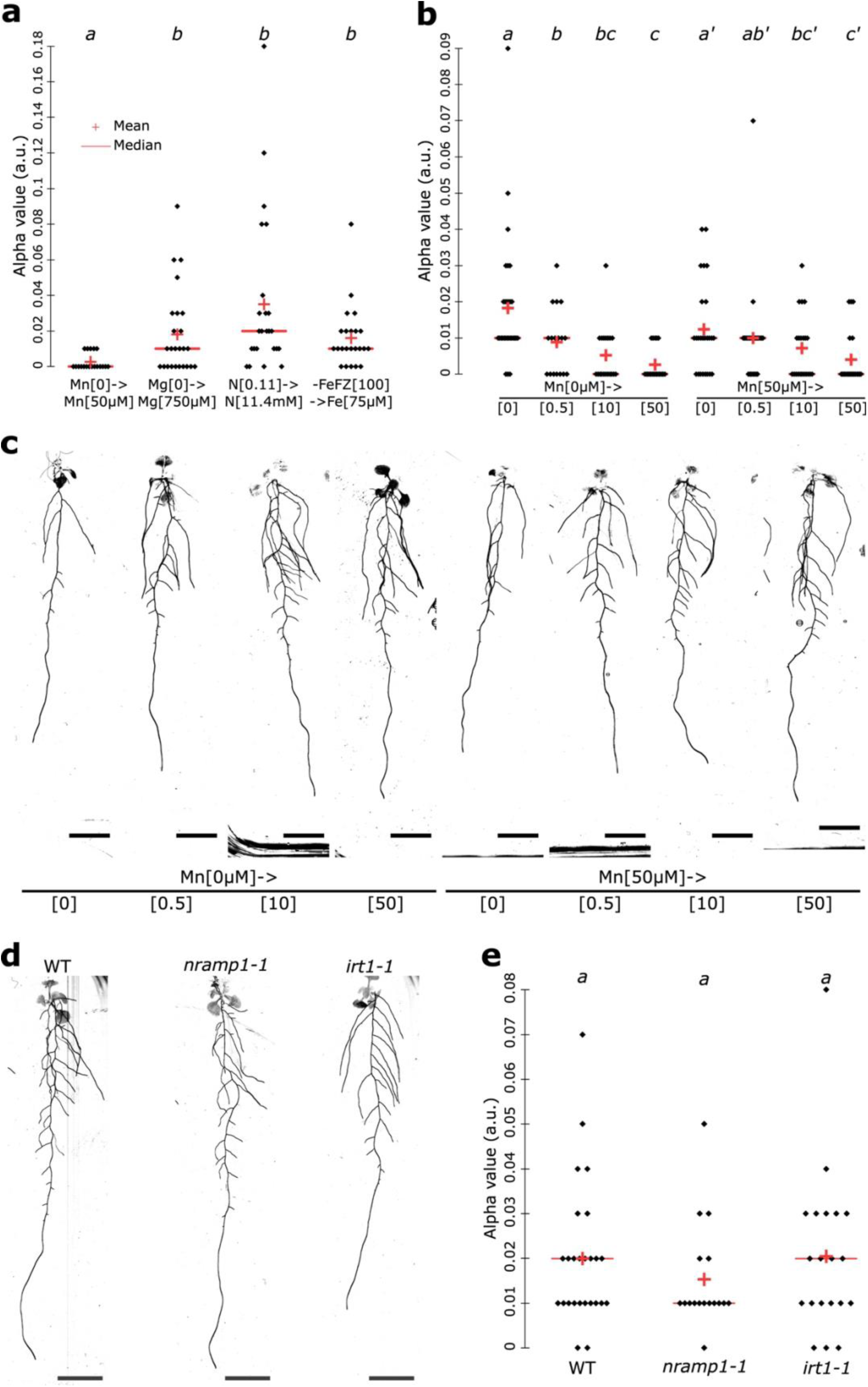
The manganese levels fine tune the RSA cost-efficiency tradeoff in a concentration-dependent manner. **a**, Quantification of the alpha value for 17-day-old WT plants grown for 5 days in the lowest levels of nutrients (Mn 0µM, N 0,11µM and Mg 0µM) and then transferred to the highest levels of nutrients (Mn 50µM, N 11.4µM and Mg 750µM) for 12 days. **b-c**, Quantification of the alpha value for 17-day-old WT plants grown for 5 days on agar plates under 0µM or 50µM then transferred to agar plates containing 4 concentrations of Mn, 0µM, 0.5µM, 10µM and 50µM (**b**) and the representative images (**c**). [two-way Kruskal-Wallis coupled with post hoc Steel-Dwass-Critchlow-Fligner procedure was performed, letters indicate statistical differences (p < 0.05)]. **d-e**, Representative images of the indicated genotypes (**d**) and the corresponding quantification of the alpha value (**e**). [two-way ANOVA, Tukey HSD test, letters indicate statistical differences (p < 0.05)]. The red crosses depict the mean and the red bars the median. Scale bar, 1cm.

## CONCLUSION

The present study provides compelling evidence that the root system architecture of Arabidopsis, much like that of *Solanum pimpinellifolium*, consistently adheres to Pareto optimality in balancing the tradeoff between transport efficiency and growth cost. Using the newly developed Ariadne software, which enables robust, semi-automated analysis of RSA topology, we demonstrated that Arabidopsis root systems grown under both natural and controlled conditions are significantly closer to the Pareto front than randomly generated root architectures. This finding underscores that real plant root systems are not random but rather are developmentally and genetically programmed to achieve an optimal compromise between efficient resource transport and minimal construction cost.

The alpha values derived from these analyses further reveal that Arabidopsis RSAs are biased towards maximizing transport efficiency, even at some expense to growth cost. This bias is evident in both adult and juvenile stages, but the exact position on the Pareto front shifts with development, suggesting a dynamic, genetically encoded regulation of RSA optimality. We also showed the influence of environmental factors, such as water availability (osmotic stress), on the cost-efficiency tradeoff, as reflected by changes in alpha values. The application of machine learning approaches, particularly the Discovery Engine in this study, allowed for high-throughput, interpretable identification of key genetic and environmental regulators of RSA traits. This approach confirmed the importance of developmental stage, genotype (notably the *hy5;chl1-5* mutant), and manganese levels in modulating the tradeoff between growth cost and transport efficiency. These results collectively suggest that RSA plasticity is governed by a genetically constrained framework that is responsive to environmental cues, enabling plants to maintain near-optimal function under diverse conditions.

In summary, our results demonstrate that the root architectures of Arabidopsis like those of *Solanum pimpinellifolium* are consistently organized according to Pareto optimal principles. This organization is not static; rather, it is developmentally regulated and environmentally responsive, allowing plants to adjust their RSA to maintain an optimal balance between the competing demands of efficient resource transport and minimal construction cost. The ability to maintain Pareto optimality across a range of conditions highlights the evolutionary advantage of this strategy, likely contributing to plant resilience, resource use efficiency, and potential for carbon sequestration. Understanding the genetic and physiological bases of this Pareto optimality will be crucial, as it will enable targeted interventions to enhance crop performance and adaptability in the face of environmental challenges.

## Supporting information

Mat & Met

## ACKNOWLEDGEMENTS

We thank all Busch lab members for critical and fruitful discussions. This study was funded by the National Institute of General Medical Sciences of the National Institutes of Health (grant number R01GM127759) and funds from the Salk Institute for Biological Studies to W.B. It was further supported by gifts to the Salk Institute’s Harnessing Plants Initiative (HPI) from the Bezos Earth Fund, the Hess Corporation, and through the TED Audacious Project. M.P.P was supported by a long-term postdoctoral fellowship (LT000340/2019L) by the Human Frontier Science Program Organization.

## AUTHOR CONTRIBUTIONS

M.P.P and W.B. conceived the study. M.P.P. was responsible for all experiments and designed all experiments described in the manuscript. K.F. designed and developed the Ariadne software and performed preliminary results for RSA screen in multiple different nutritional conditions. P.M., A.M, T.H., J.S., and A.R., traced the RSA using Ariadne. E.B. design the pipy package and is in charge of keeping Ariadne software up to date. L.B., genotyped and took care of the plants. C.C. perform the RSA experiment under sorbitol. A.C., provide guidance and shared the Pareto optimal script. C.P.., developed the script to read RSML files. L.M.R., R.M. and A.T. all contributed to data preprocessing, modelling, extraction of insights and reporting the results using the Discovery Engine, whilst J.R. supervised these efforts. S.N., A.M., S.L., M.P.P, and W.B. provided funding, resources, and supervision. M.P.P. and W.B. wrote the manuscript, and all the authors discussed the results and commented on the manuscript.

## Code availability

https://github.com/Salk-Harnessing-Plants-Initiative/Ariadne

---

**Figure S1:**
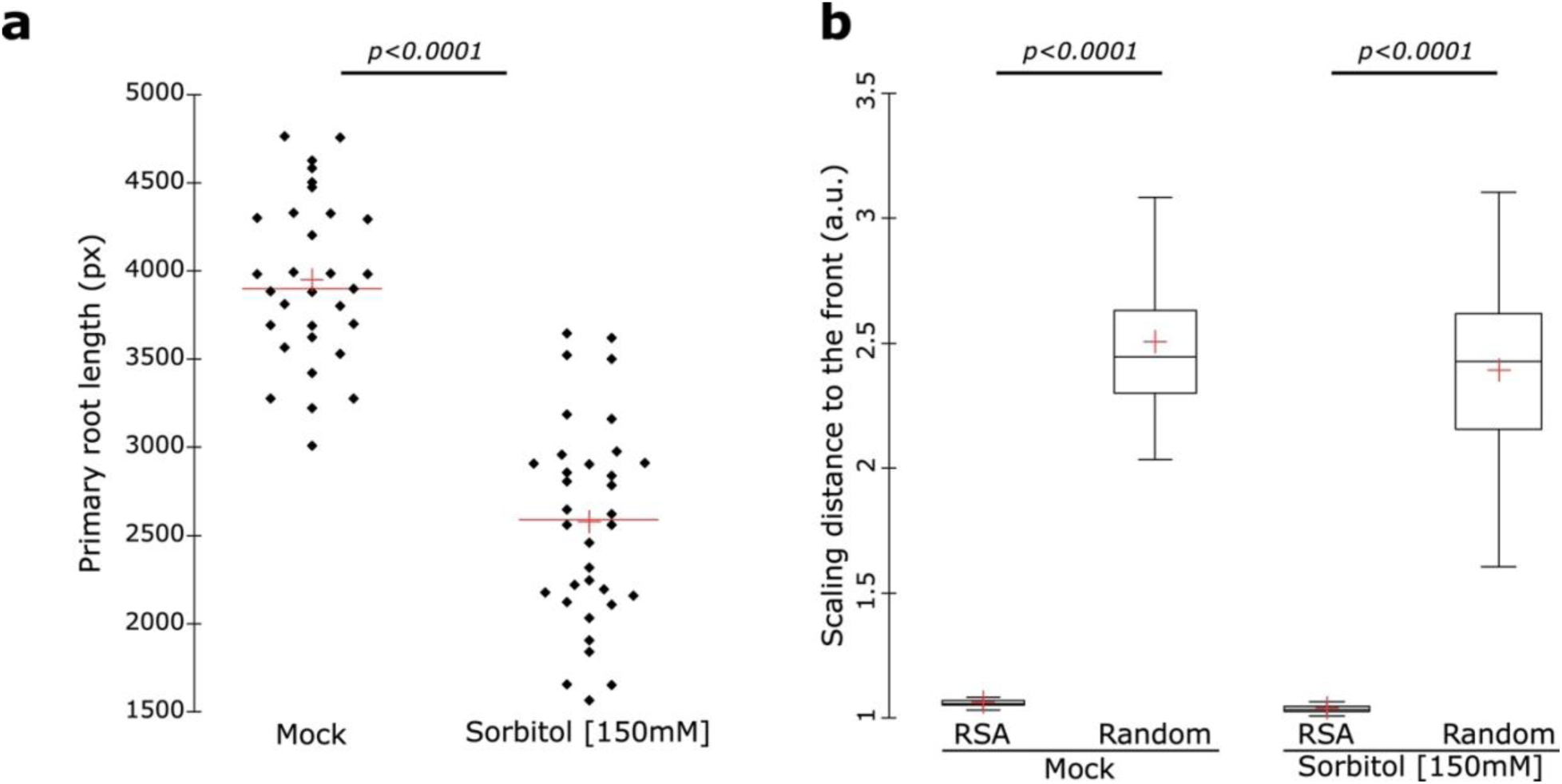
Osmotic stress induces a root growth decrease. **a**, graph depicting the primary root length of plants grown under control and sorbitol at 150mM. [Two-ways Student t-test, (p=0.05)]. **b**, graph depicting the scaling distance to the front in mock and Sorbitol conditions for RSA and random architectures. [Two-ways Student t-test (p=0.05)]. The red crosses depict the mean and the red bars the median.

**Figure S2:**
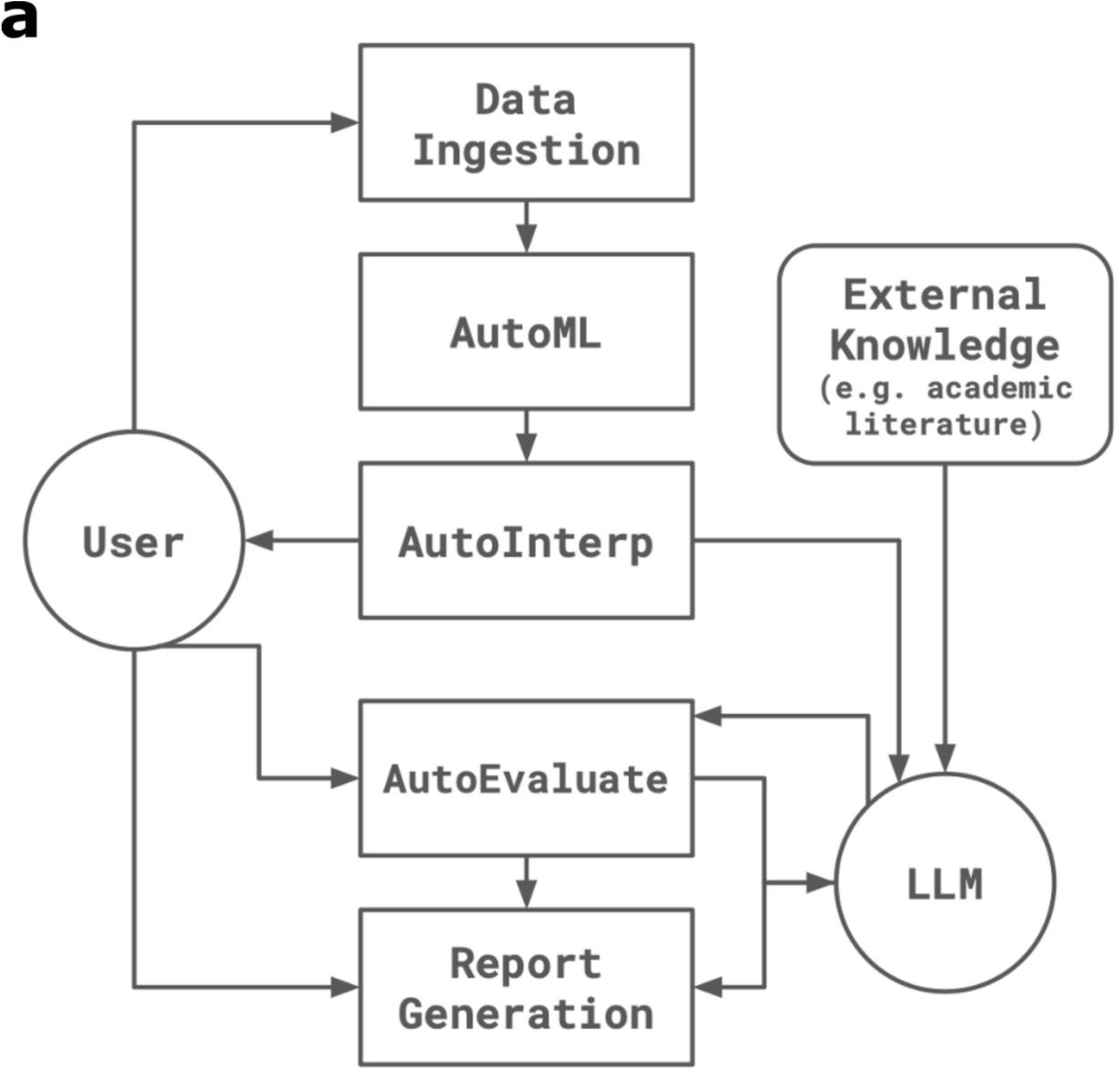
Schematic of the Discovery Engine. a, Schematic representation of the Discovery Engine and the different steps

**Figure S3:**
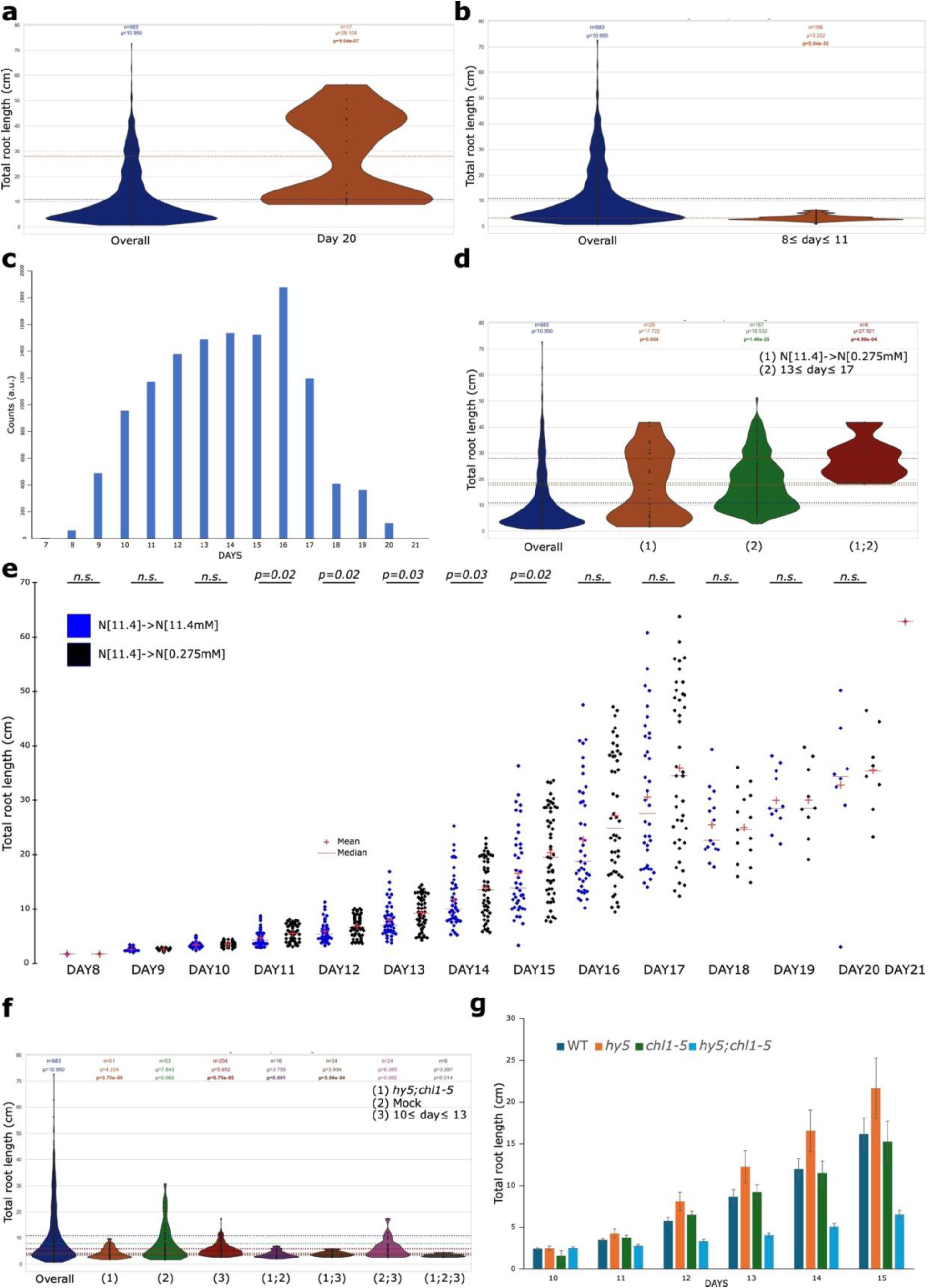
Analysis of the raw data for the total root length. **a**, Quantification of the total root length throughout the entire dataset (Overall) and at day 20. **b**, Quantification of the total root length throughout the entire dataset (Overall) and between day 8 to 11. **c,** Quantification of the number of counts per day in the entire dataset. **d**, Quantification of the total root length throughout the entire dataset (Overall), for plants grown for 5 days on agar plates under 11.4mM of N then transferred to agar plates containing 0.275mM of N and for plants between day 8 to 11 and their different combinations. **e,** Quantification of the total root length for plants grown for 5 days on agar plates under 11.4mM of N then transferred to agar plates containing 0.275mM or 11.4mM of N per day. **f**, Quantification of the total root length throughout the entire dataset (Overall), for the *hy5;chl5-1* mutant, for plants grown under mock conditions, for plants between day 10 to 13 and their different combinations. **g**, Quantification of the total root length per day for the indicted genotypes in mock conditions. [Two-ways Student t-test, (p=0.05)]. n, depicts the number of samples, µ, the mean and p, the p-value compared the overall group. The red crosses depict the mean and the red bars the median.

**Figure S4:**
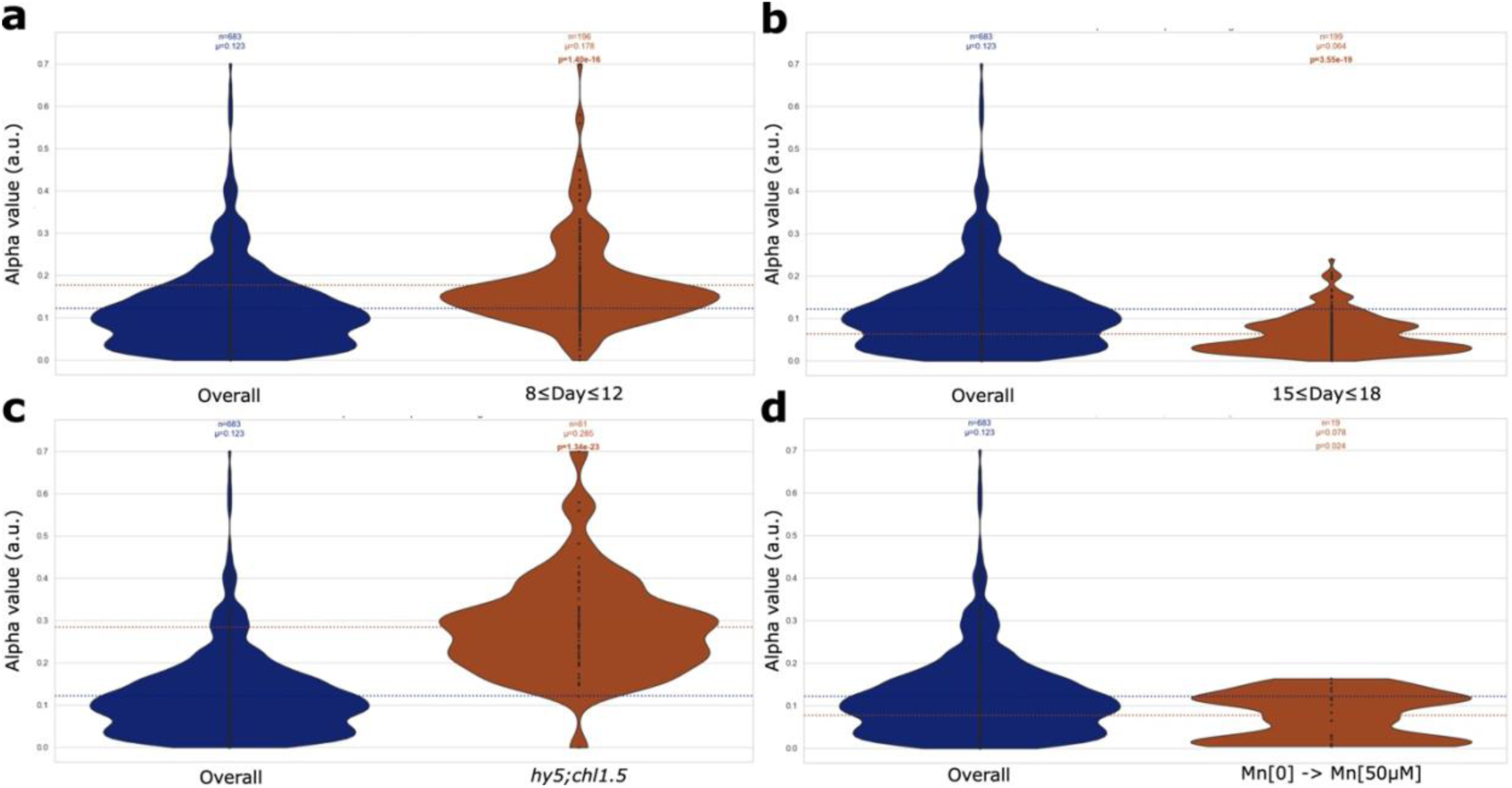
Analysis of the raw data for the alpha value. **a**, Quantification of the alpha value throughout the entire dataset (Overall) and between day 8 to 12. **b**, Quantification of the alpha value throughout the entire dataset (Overall) and between day 15 to 18. **c**, Quantification of the alpha value throughout the entire dataset (Overall) or for the *hy5;chl5-1* mutant. **d**, Quantification of the alpha value throughout the entire dataset (Overall), for plants grown for 5 days on agar plates under 0µM of Mn then transferred to agar plates containing 50µM of Mn. [Two-ways Student t-test, (p=0.05)]. n, depicts the number of samples, µ, the mean and p, the p-value compared the overall group.

**Figure S5:**
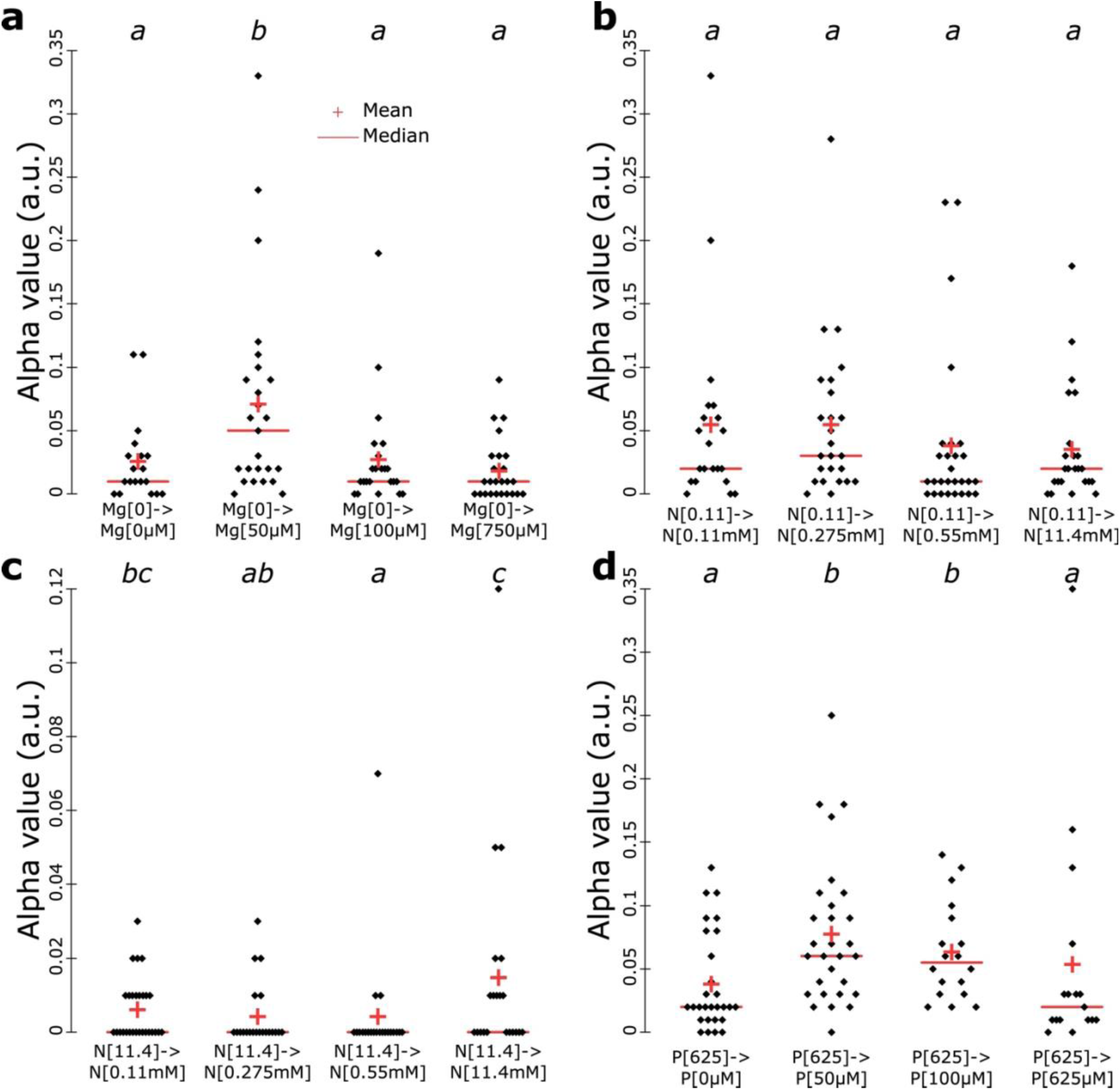
The Mg, P and N does not modulate the alpha value in a concentration-dependent manner. **a**, Quantification of the alpha value for 17-day-old plants grown for 5 days on agar plates under 0µM of Mg then transferred to agar plates containing 0, 50, 100 and 750µM of Mg. **b**, Quantification of the alpha value for 17-day-old plants grown for 5 days on agar plates under 0.11mM of N then transferred to agar plates containing 0.11, 0.275, 0.550 and 11.4mM of N. **c**, Quantification of the alpha value for 17-day-old plants grown for 5 days on agar plates under 11.4mM of N then transferred to agar plates containing 0.11, 0.275, 0.550 and 11.4mM of N. **d**, Quantification of the alpha value for 17-day-old plants grown for 5 days on agar plates under 625µM of P then transferred to agar plates containing 0, 50, 100 and 625µM of P. [two-way Kruskal-Wallis coupled with post hoc Steel-Dwass-Critchlow-Fligner procedure was performed, letters indicate statistical differences (p < 0.05)]. The red crosses depict the mean and the red bars the median.

**Table S1: Root system architectural traits extracted by Ariadne**

**Table S2: Nutritional growth conditions used for the large-scale screens**

**Table S3: Entire dataset of the RSA**

**Table S4: Results of Discovery Engine for the total root growth and the alpha value**

**Supplemental 1: Evaluation of the trained models**

