## Supplementary material for "Growth Cost and Transport Efficiency Tradeoffs Define Root System Optimization Across Varying Developmental Stages and Environments in Arabidopsis": Mat & Met

### MATERIAL & METHODS

**Plant materials and growth conditions.** For surface sterilization, *Arabidopsis thaliana* seeds that had been produced under uniform growth conditions and were placed for 1 h in opened 1.5-mL Eppendorf tubes in a sealed box containing chlorine gas generated from 130 mL of 10% sodium hypochlorite and 3.5 mL of 37% hydrochloric acid. For stratification, seeds were imbibed in water and stratified in the dark at 4 °C for 2-3 days. Seeds were then put on the surface agar media using 12-cm x 12-cm square plates. The standard media for gradient of Fe, Mg, Mn, N and P contain, 750 µM of $MgSO_4$-7H2O, 625 µM of $KH_2PO_4$, 1000 µM of $NH_4NO_3$, 9400 µM of $KNO_3$, 1500 µM of $CaCL_2$-$2H_2O$, 0.055 µM of $CoCL_2$-$6H_2O$, 0.053 µM of $CuCl_2$-$2H_2O$, 50 µM of $H_3BO_3$, 2.5 µM of KI, 50 µM of $MnCl_2$-$4H_2O$, 0.52 µM of $Na_2MoO_4$-$2H_2O$, 15 µM of $ZnCl_2$, 75 µM of Na-Fe-EDTA, 1000 µM of MES adjusted to pH 5.5 with KOH[1]. For the medias with gradient levels of Fe, Mg, Mn, N and P, please refer to Table S2. For the media preparation of N0, P0 and N0P0, MS half media (Caisson lab, Cat. MSP33) Nitrogen-deficient media (Caisson lab, Cat. MSP19), or Phosphate deficient media (Caisson lab, Cat. MSP21) were used with pH 5.7, 0.8% micropropagation Type 1 phytoagar (Caisson), and supplement of nitrogen or phosphate source according to the previous report (Gruber) (N550: 50µM NH4NO3,450µM KNO3, and 8900µM KCl, P125: 100µM KH2PO4 and 525µMKCl)[2]. The media supplemented with hormones or chemical were half MS media with 1% sucrose and 1% phytagel agar. The media used to perform sorbitol treatment contained *$KH_2PO_4$* 1 mM, $MgSO_4$ 1 mM, $K_2SO_4$ 250 *µ*M, $CaCl_2$ 250 *µ*M, Na-Fe-EDTA 100 *µ*M, $KNO_3$ 10 mM, KCl 50 *µ*M, $H_3BO_3$ 30 *µ*M, $MnSO_4$ 5 *µ*M, $ZnSO_4$ 1 *µ*M, $CuSO_4$ 1 *µ*M, $(NH_4)_6Mo_70_{24}$ 0,1 *µ*M[3]. The, *brl3-2* (*SALK_006024C)*[4], *chl1-5*[5], *hy5-215*[6], *hy5-215;chl1-5*[2], *nramp1-1*[7] and *irt1-1*[8] are in Col-0 background and were previously described and characterized. Plants were grown in long day conditions (16/8h) in walk in growth chambers at 21°C, 100*µ*M light intensity, 60% humidity. During the nighttime, temperature was decreased to 15°C.

#### Chemical treatment with sorbitol

Seeds were sowed in the media described in 'Plant material and conditions' and stratified for 2-3 days at 4°C1. 6-day-old seedlings were transferred to plates containing the same media and supplemented with 150mM of Sorbitol (CAS 50-70-4, CAT#1876).

#### INTERPRETABLE MACHINE LEARNING: DISCOVERY ENGINE

The Discovery Engine was deployed on the Ariadne output as described here [9]. Ariadne data was cleaned and preprocessed before training the neural network as follows:

• Merged accidental duplicate categorical values in the "Nutrient" column, where 'brl3-2' and 'brl3_2' referred to the same nutrient but were labeled inconsistently.

• Duplication handling: For every target we can merge duplicate input feature rows to create a final dataset of size 683 rows.

#### QUANTIFICATION

**Root system architecture traits.** Seeds were sowed in the indicated media described in Table S2 or in the section "Plant materials and growth conditions". They were stratified for 2–3days at 4°C. Five days after planting, 5 plants were transferred to 12×12-cm plates. After transfer, the plates were scanned using CCD flatbed scanner every 24 h until the root tip of one of the plant reach the bottom of the plate as previously described[10]. The images were analyzed using Ariadne software. The user opens the image using the "Trace" function and proceeds to manual tracing of the root system day after day or at the required day. A JSON file is created and then analyze using the "Analyze" function. The results are displayed on a .csv file and contain 18 different RSA traits listed in the Table S1. The source code is provided here (https://github.com/Salk-Harnessing-Plants-Initiative/Ariadne) And the software is fully available here (https://pypi.org/project/ariadne-roots/).

**STATISTICS & GRAPHS**

Each experiment has been repeated independently at least twice, as in every case the same trend has been recorded for independent experiments and the data have been pooled for further statistical analysis. Each sample were subjected to four different normality tests (Jarque-Bera, Lilliefors, Anderson-Darling and Shapiro-Wilk), samples were considered as a Gaussian distribution when at least one test was significant ($p<0.05$) using Xlstat.

- As a normal distribution was observed a one or two-ways ANOVA coupled with post hoc Tukey test was performed ($p<0.05$) using Xlstat.

- As a normal distribution was observed a one or two-ways ANOVA coupled with post hoc Fisher test was performed ($p<0.05$) using Xlstat:

- As a normal distribution was observed an independent two-ways student test was performed ($p=0.05$) using Xlstat.

- As a normal distribution was not observed at two-way Kruskal-Wallis coupled with post hoc Steel-Dwass-Critchlow-Fligner procedure was performed ($p<0.05$) using Xlstat.

The type of test performed in indicated in the figure legend.
In each graph, according to the box plot representation it depicts, the minimum as the lowest data point in the data set excluding any outliers, the maximum as the maximum as the lowest data point in the data set excluding any outliers, the median as the mid value in the data set, the first quartile as the median of the lower half of the data set, and the third quartile as the median of the upper half of the data set. The cross represents the mean of the data set. Cercles represent each individual measurements composing the data set.
