## Supplemental_S1 for "Growth Cost and Transport Efficiency Tradeoffs Define Root System Optimization Across Varying Developmental Stages and Environments in Arabidopsis"

Evaluation of the Trained Models

For all models we use Mean Squared Error (MSELoss) as the loss function during training. For evaluation, we compute Mean Absolute Error (MAE), Root Mean Squared Error (RMSE) and R-squared (R^2^) score.

1. Total Root Length

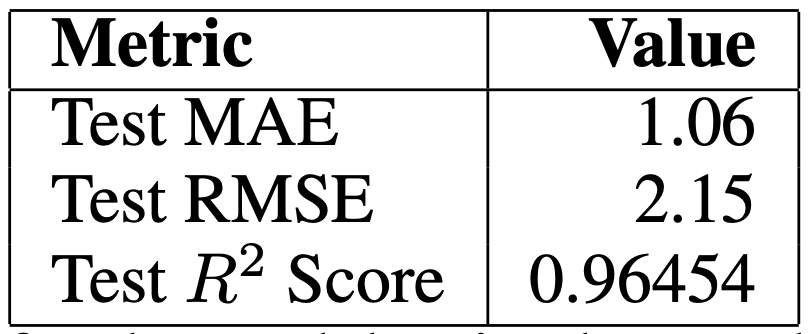

Table 1: Test metrics for the model trained to predict Total Root Length

2. Alpha

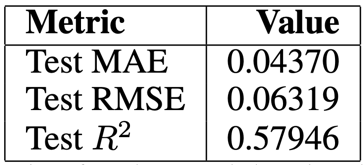

Table 2: Test metrics for the model trained to predict alpha
